# Extensive In Silico Analysis of *ATL1* Gene: Discovered Five Mutations that may Cause Hereditary Spastic Paraplegia Type 3A

**DOI:** 10.1101/818302

**Authors:** Mujahed I. Mustafa, Naseem S. Murshed, Abdelrahman H. Abdelmoneim, Miysaa I. Abdelmageed, Nafisa M. Elfadol, Abdelrafie M. Makhawi

## Abstract

**BACKGROUND:** Hereditary spastic paraplegia type 3A (SPG3A) is a neurodegenerative disease inherited type of Hereditary spastic paraplegia (HSP). It is the second most frequent type of HSP; which Characterized by muscle stiffness with paraplegia and early-onset of symptoms. This is the first translational bioinformatics analysis in a coding region of *ATL1* gene which aims to categorize nsSNPs to be used as genomic biomarkers; also it may play a key role in pharmacogenomics by evaluating drug response for this disabling disease.

**METHODS:** The raw data of *ATL1* gene were retrieved from dbSNP database, and then run into numerous computational analysis tools. Additionally; we submitted the common six deleterious outcomes from the previous functional analysis tools to I-mutant 3.0, and MUPro respectively, to investigate their effect on structural level. The 3D structure of *ATL1* was predicted by RaptorX and modeled using UCSF Chimera to compare the differences between the native and the mutant amino acids.

**RESULTS:** Five nsSNPs out of 249 were classified as the most deleterious (rs746927118, rs979765709, rs119476049, rs864622269, rs1242753115).

**CONCLUSIONS:** In this study the impact of nsSNPs in the *ATL1 gene* was investigated by various bioinformatics tools, that revealed five nsSNPs (V67F, T120I, R217Q, R495W and G504E) are deleterious SNPs, which have a functional impact on ATL1 protein; and therefore, can be used as genomic biomarkers specifically before 4 years old; also it may play a key role in pharmacogenomics by evaluating drug response for this disabling disease.

## 1. Introduction

Hereditary spastic paraplegia type 3A (SPG3A) is a neurodegenerative disease inherited type of Hereditary spastic paraplegia (HSP) [1] SPG3A is the second most frequent type of HSP.[2-4] SPG3A Characterized by muscle stiffness with paraplegia and early-onset of symptoms; [5, 6] however, in rare cases extremely severe characteristic with neonatal onset have been reported.[6] SPG3A had been recognized in different populations.[7-10]

Insight into the molecular basis of HSP is increasing rapidly; different genes cause clinically indistinguishable HSP types.[11-13] SPG3A is triggered by heterozygous mutations in *ATL1* gene (14q22.1), but the genotype-phenotype correlation still unknown. [14-17] Mutations in the *ATL1* gene are more common with early-onset HSP patients.[18] These mutations probably cause atypical activity of atlastin-1 protein, which damages the transmission of neurons.[4] The population databases (http://exac.broadinstitute.org/) claims a rare benign SNPs also found in *ATL1* in healthy persons. Besides, a rare mutations in the *ATL1* have been reported in axonal motor neuropathy [17, 19] and hereditary sensory neuropathy type I patients.[20] Most importantly, given the absence of cure, it’s vital for appropriate genetic counseling for the manifestation of any variants related to SPG3A, because the treatment is symptomatic; muscle stiffness can be treated with oral baclofen or tizanidine; Physical therapy should be combined to improve the life quality of the patient.[1]

The problematic issue stems from the point that the effects of genetic differences on macromolecules function vary extensively making it difficult to decode genotype-phenotype correlation; some study suggest that linkage analysis is a beneficial approach to assist in understanding this correlation.[21]

This is the first translational bioinformatics analysis in coding region of *ATL1* gene which aims to categorize nsSNPs could use as genomic biomarkers specifically before 4 years old; also it may play a key role in pharmacogenomics by evaluating drug response for this disabling disease.[22-27]

## 2. Methods

### 2.1 Data Mining

The data of *ATL1* gene were retrieved from dbSNP database, (http://www.ncbi.nlm.nih.gov/snp/); while the protein reference sequence (Q8WXF7) was obtained from UniProt database (https://www.uniprot.org/).

### 2.2 Functional analysis

#### 2.2.1 SIFT

SIFT was used to observe the effects of each amino acid substitution on protein function. SIFT predicts damaging SNPs on the basis of the degree of conserved amino acid residues in aligned sequences to the closely related sequences, gathered through PSI-BLAST.[28]

#### 2.2.2 PolyPhen-2

PolyPhen 2 stands for polymorphism phenotyping version 2. We used PolyPhen to study potential effects of each amino acid substitution on structural and functional properties of the protein by considering physical and comparative approaches. The input data needs accession number, position of mutations, native and altered amino acids.[29]

#### 2.2.3 PROVEAN

It predicts whether an amino acid substitution has an effect on the biological function of a protein grounded on the alignment-based score. If the PROVEAN score ≤-2.5, the protein variant is predicted to have a “deleterious” effect, while if the PROVEAN score is >-2.5, the variant is predicted to have a “neutral” effect.[30]

#### 2.2.4 SNAP2

It is a trained functional analysis web-based tool that differentiates between effect and neutral SNPs by taking a variety of features into account. SNAP2 got accuracy (effect/neutral) of 83%. It is considered an important and substantial enhancement over other methods.[31]

#### 2.2.5 SNPs&GO

It is a support vector machine (SVM) based on the method to accurately predict the disease related mutations from protein sequence. The probability score higher than 0.5 reveals the disease related effect of mutation on the parent protein function.[32]

#### 2.2.6 PHD-SNP

It is an online Support Vector Machine (SVM) based classifier, is optimized to predict if a given single point protein mutation can be classified as disease-related or as a neutral polymorphism.[33]

### 2.3 Stability analysis

#### 2.3.1 I-Mutant 3.0

Change in protein stability disturbs both protein structure and protein function. I-Mutant is a suite of support vector machine. It offers the opportunity to predict the protein stability changes upon single-site mutations. The FASTA format sequence of ATL1 protein was retrieved from UniProt that used as an input to predict the mutational effect on protein and stability RI value (reliability index) computed.[34]

#### 2.3.2 MUPro

It is a support vector machine-based tool for the prediction of protein stability changes upon nonsynonymous SNPs. The value of the energy change is predicted, and a confidence score between −1 and 1 for measuring the confidence of the prediction is calculated. A score <0 means the variant decreases the protein stability; conversely, a score >0 means the variant increases the protein stability.[35]

### 2.4 Biophysical and visualization analysis

#### 2.4.1 Project HOPE

It is a server to search structural data from a several database such as UniProt. The FASTA format sequence of ATL1 protein was retrieved from UniProt that used as an input to predict the biophysical validation for our SNPs of interest. The main aims for the submissions in Project HOPE are to analysis and confirm results that we had it earlier.[36]

#### 2.4.2 RaptorX

The full 3D structure of human ATL1 protein is not available in the Protein Data Bank. Hence, we used RaptorX to generate a 3D structural model for wild-type ATL1. The FASTA format sequence of XK protein was retrieved from UniProt that used as an input to predict the 3D structure of human ATL1 protein.[37]

#### 2.4.3 UCSF Chimera

UCSF Chimera is a highly extensible program for interactive visualization and analysis of molecular structures and related data, including density maps, supramolecular assemblies, sequence alignments, docking results, conformational analysis Chimera (version 1.8).[38]

### 2.4 ConSurf server

It is a web server suggests evolutionary conservation reviews for proteins of known structure in the PDB. ConSurf detect the similar amino acid sequences and run multi alignment approaches. The conserved amino acid across species detects its position using specific algorisms.[39]

### 2.5 GeneMANIA

It is web server creating proposition about gene function, investigating gene lists and prioritizing genes for functional assays. The high accuracy of the GeneMANIA prediction algorithm and large database make the GeneMANIA a useful tool for any biologist.[40]

### 2.6 ClinVar

It is a public archive of reported studies of the relationships among human variations and phenotypes, with supporting evidence. We used it to compare our prediction approach with the clinical one.[41]

### 2.7 Variant Effect Predictor (VEP)

The Ensembl Variant Effect Predictor software provides toolsets for an organized approach to annotate and assist for prioritization of mutations. The input data format was a list of variant identifiers while the output was filtered by choosing 1000 genome combined population to expend the population coverage.[42]

## 3. Results

The total number of SNPs in the coding region that recovered from NCBI was 249 nsSNPs, and these SNPs were submitted into different functional analysis online tools, (figure 1) There are 98 out of 249 nsSNPs were found to be affected by SIFT, 118 damaging SNPs (44 possibly damaging & 74 probably damaging) by Pholyphen-2 and 122 were found to be deleterious by PROVEAN; the triple-positive damaging SNPs were filtered from the earlier three online tools, out of 41 SNPs there were 8 predicted damaging by SNAP2. (Table 1) After second filtration the number of SNPs decreased to 8, then were submitted into SNPs&GO and PhD-SNP and P-Mut, to give more accurate results on their effect on the functional impact; the triple positive in the three tools was five SNPs (Table 2); on the other hand, the stability analysis on these five SNPs were tested by I-Mutant3.0 and MUPro, the stability analysis revealed that, all five SNPs decrease the protein stability, except for one SNP (T120I) that was predicted by I-Mutant3.0 to increase protein stability.(Table 3).

**Table 1:** Pathogenic nsSNPs predicted by different online tools:

| dbSNP rs# | SUB | Sift Prediction | Score | Polyphen prediction | Score | PROVEAN prediction | Score | SNAP2 Prediction | Score |
| --- | --- | --- | --- | --- | --- | --- | --- | --- | --- |
| rs778130435 | F46S | Affect | 0 | probably damaging | 1 | Deleterious | -7.014 | Effect | 48 |
| rs746927118 | V67F | Affect | 0 | probably damaging | 1 | Deleterious | -4.221 | Effect | 36 |
| rs979765709 | T120I | Affect | 0 | probably damaging | 1 | Deleterious | -5.216 | Effect | 83 |
| rs149031604 | T147I | Affect | 0 | probably damaging | 1 | Deleterious | -5.544 | Effect | 79 |
| rs119476049 | R217Q | Affect | 0 | probably damaging | 1 | Deleterious | -3.749 | Effect | 95 |
| rs1212638776 | A359T | Affect | 0 | probably damaging | 1 | Deleterious | -3.722 | Effect | 20 |
| rs864622269 | R495W | Affect | 0 | probably damaging | 1 | Deleterious | -7.265 | Effect | 73 |
| rs1242753115 | G504E | Affect | 0 | probably damaging | 1 | Deleterious | -7.659 | Effect | 78 |

**Table 2:** The most damaging SNPs predicted by different online tools:

| dbSNP rs# | SUB | SNPs&GO Prediction | RI | Probability | PHD-SNP Prediction | RI | Probability | P-Mut Prediction | Score |
| --- | --- | --- | --- | --- | --- | --- | --- | --- | --- |
| rs746927118 | V67F | Disease | 1 | 0.559 | Disease | 7 | 0.827 | Disease | 0.88 (92%) |
| rs979765709 | T120I | Disease | 2 | 0.622 | Disease | 5 | 0.765 | Disease | 0.92 (94%) |
| rs119476049 | R217Q | Disease | 3 | 0.626 | Disease | 6 | 0.81 | Disease | 0.84 (90%) |
| rs864622269 | R495W | Disease | 5 | 0.744 | Disease | 8 | 0.921 | Disease | 0.79 (89%) |
| rs1242753115 | G504E | Disease | 1 | 0.557 | Disease | 7 | 0.85 | Disease | 0.73 (87%) |

**Table 3:**
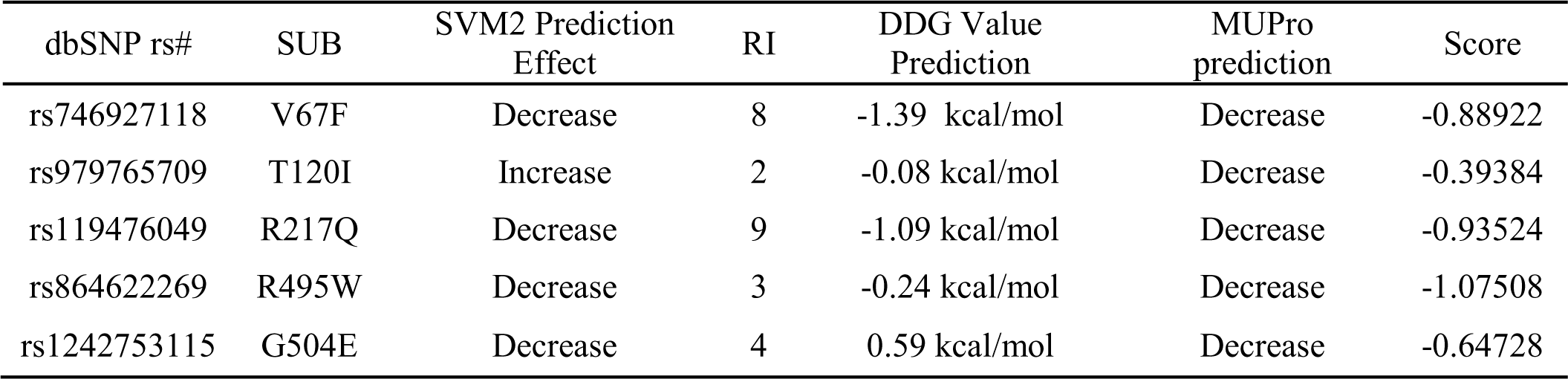
Protein structural stability calculated using I-Mutant 3.0 and MUPro:

**Figure 1:**
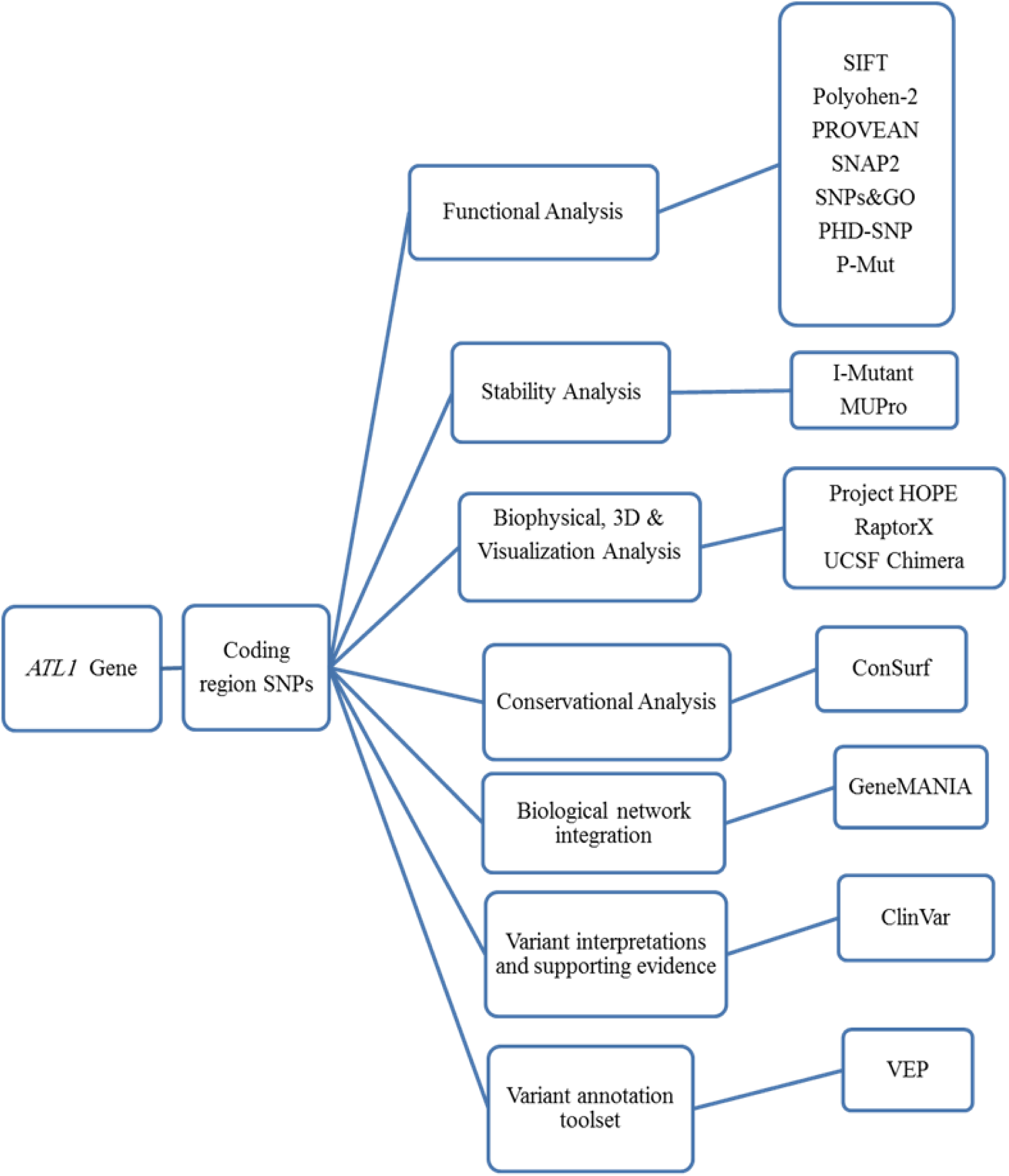
Graphic demonstration of *ATL1* gene work flow.

**Figure 2:**
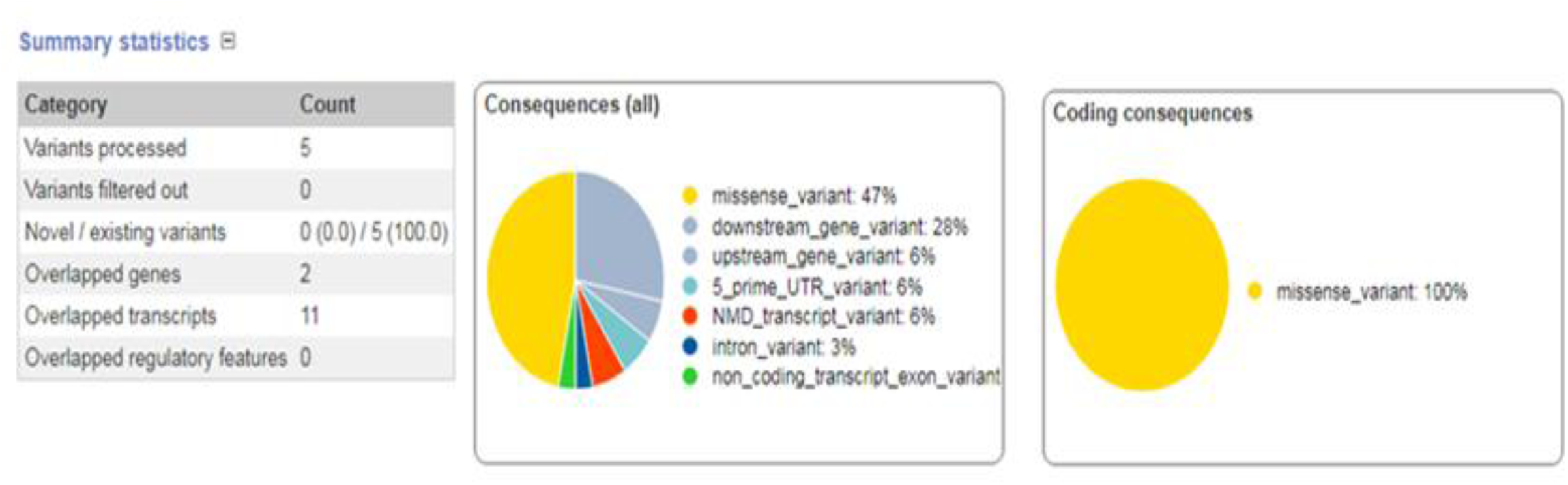
Summary pie charts and statistics.

**Figure 3:**
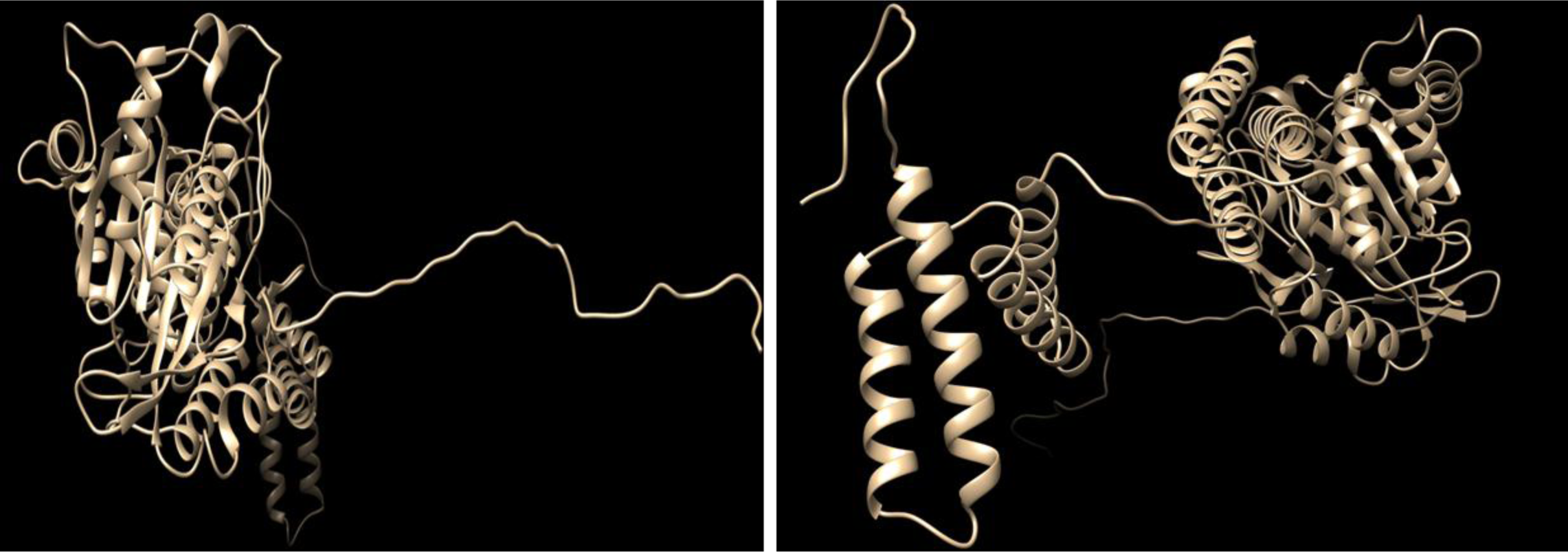
The 3D structure of ATL1 protein model by two angles.

## 4. Discussion

In vitro approach is consuming a lot of time and fees which may or may not have a positive result from the study, in the other hand, computational approach is entirely different, it saves time, with low cost and gives rapid results to improve our understanding of how variants could interrupt the protein structure and function.[43, 44]

Missense mutations are frequently found to arise at evolutionarily conserved regions. Those have a key role at structural and functional levels of the protein.[45-47] Therefore, our in silico analysis was devoted to the coding region of *ATL1* gene, which uncovered five disease-causing mutations that may cause SPG3A. The five deleterious SNPs come after extensive computational analysis, seven online tools (figure 1) were used to investigate the effect of each SNP on the functional impact, the reason why results are different in many times because they run by different sequence and structure-based algorithms; (Table1&2) while two online tools (I-Mutant and MUPro) were used to investigate the effect of each SNP on the stability impact, the analysis revealed that, all five SNPs decrease the protein stability, except for one SNP (T120I) that was predicted by I-Mutant3.0 to increase protein stability; (Table 3) thus proposing that these variants could destabilize the amino acid interactions causing functional deviations of protein to some point.

All these SNPs (V67F, T120I and G504E) were recovered from the dbSNP as untested and all were found to be deleterious mutations; while these SNPs (R217Q and R495W) were recovered as Pathogenic which agree with our finding. (Table 2)

In order to investigate the biophysical properties of these variants, Project HOPE server was used to serve this purpose, while UCSF Chimera was used to visualize the amino acid change; in (**Figure 4)**: (V67F): The amino acid Valine changes to Phenylalanine at position 67; The mutated residue is located in a domain that is important for binding of other molecules and in contact with residues in a domain that is important for the activity of the protein. The mutation might affect this interaction and thereby disturb signal transfer from binding domain to the activity domain. The mutation introduces an amino acid with different properties, which can disturb this domain and abolish its function; The wild-type residue is very conserved but the mutant residue is located near a highly conserved position.

**Figure 4:**
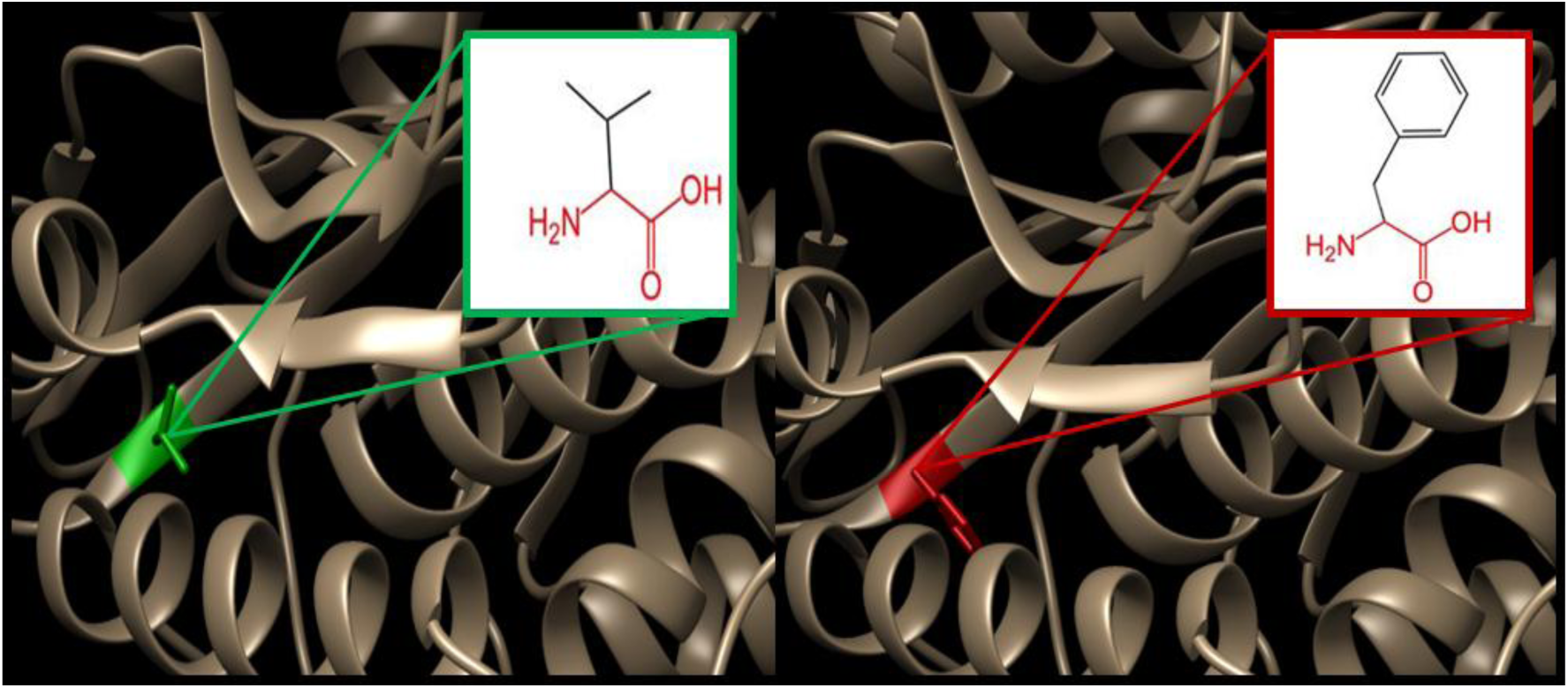
(V67F): The amino acid Valine changes to Phenylalanine at position 67

In (**Figure 5)**: (T120I): The amino acid Threonine changes to Isoleucine at position 120; The mutant residue is bigger than the wild-type residue, these differences disturb the interaction with the metal-ion (MG); These differences in properties between wild-type and mutant residue can easily cause loss of interactions with the nucleotide (GTP), which can directly affect the function of the protein. The mutation is located within a domain, annotated in UniProt as: (GB1/RHD3-type G). Only this residue type was found at this position. Mutation of a 100% conserved residue is usually damaging for the protein.

**Figure 5:**
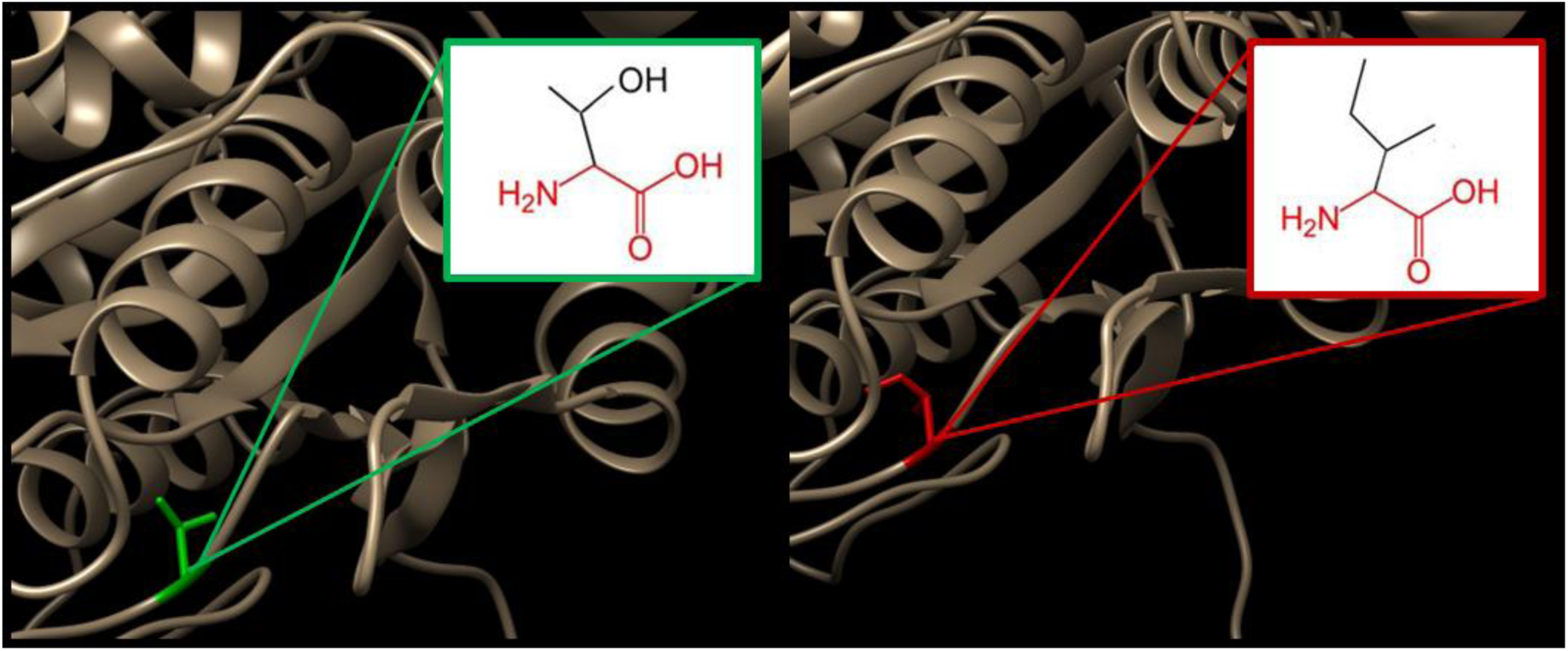
(T120I): The amino acid Threonine changes to Isoleucine at position 120.

In (**Figure 6)**: (R217Q): The amino acid Arginine changes to Glutamine at position 217; in the 3D-structure, the wild-type residue has interactions with a ligand annotated as (GDP). The difference in properties between wild-type and mutation can easily cause loss of interactions with the ligand, because ligand binding is often important for the protein’s function, this function might be disturbed by this mutation. The wild-type residue charge was positive, while the mutant residue charge is neutral; the difference in charge will disturb the ionic interaction made by the original, wild-type residue. The mutated residue is in contact with residues in another domain. It is possible that the mutation disturbs these contacts, and as such affect the protein function.

**Figure 6:**
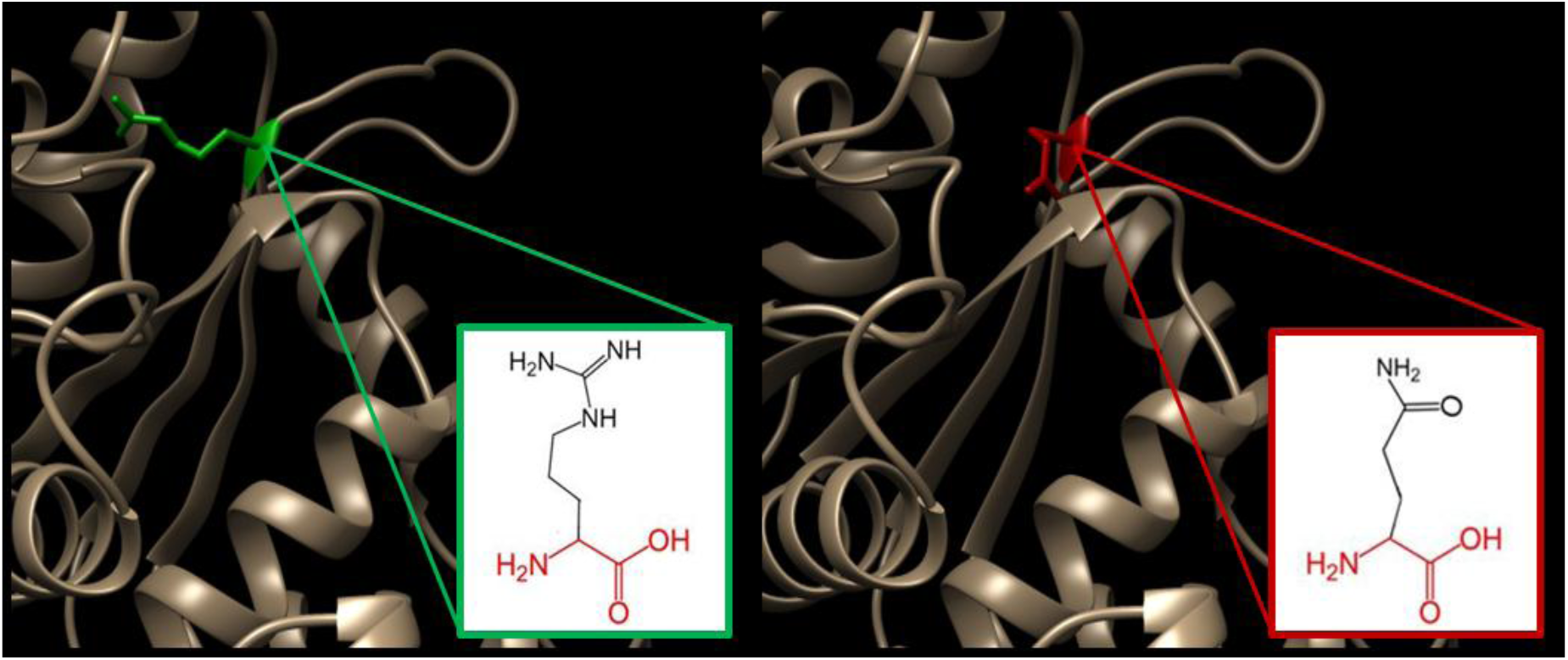
(R217Q): The amino acid Arginine changes to Glutamine at position 217.

In (**Figure 7)**: (R495W): The amino acid Arginine changes to Tryptophan at position 495; the mutation is located within a stretch of residues annotated in UniProt as a special region: Sufficient for membrane association. The differences in amino acid properties can disturb this region and disturb its function. The wild-type residue charge was positive, the mutant residue charge is neutral, and the loss of the charge can cause loss of interactions with other molecules or residues; the mutation introduces a more hydrophobic residue at this position. This can result in loss of hydrogen bonds and/or disturb correct folding.

**Figure 7:**
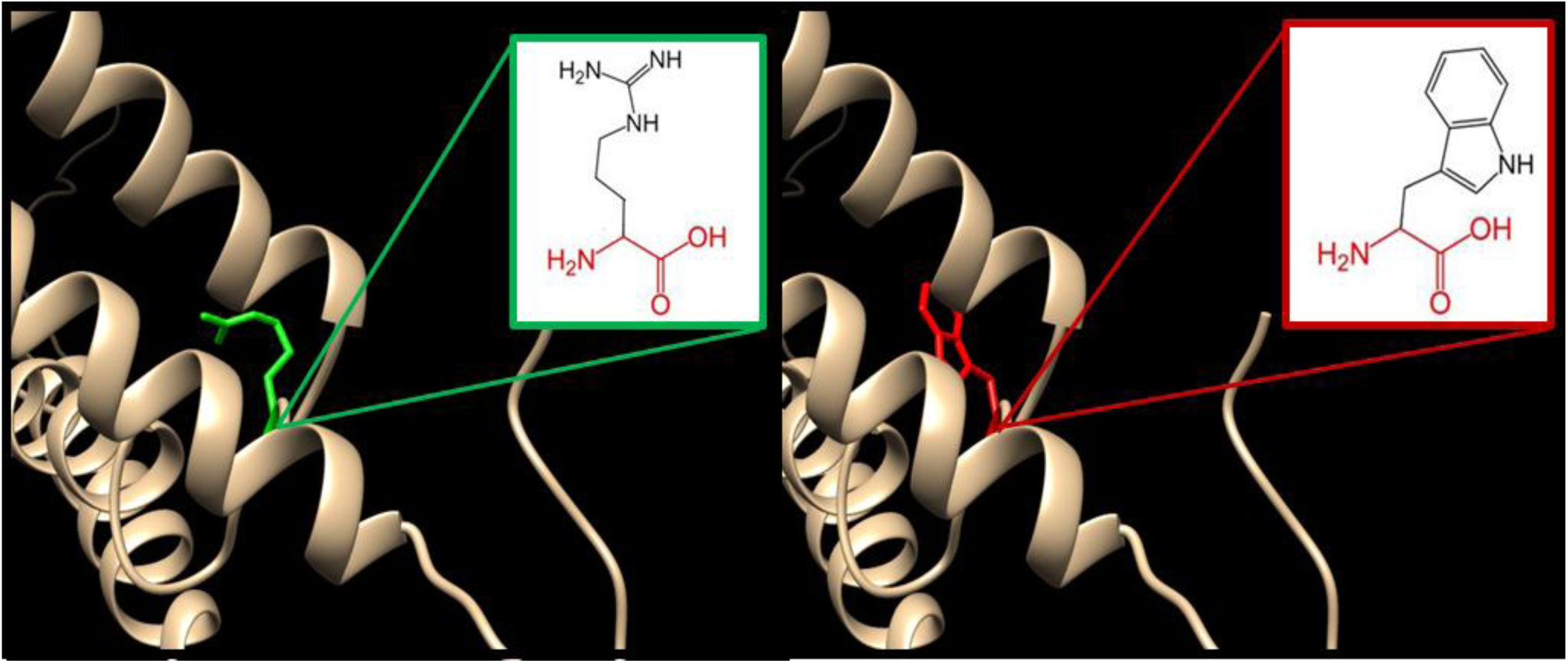
(R495W): The amino acid Arginine changes to Tryptophan at position 495.

In (**Figure 8)**: (G504E): The amino acid Glycine changes to Glutamate at position 495; the wild-type residue is a glycine, the most flexible of all residues. This flexibility might be necessary for the protein’s function. Mutation of this glycine can abolish this function. The wild-type residue charge was neutral, the mutant residue charge is negative, and this can cause repulsion of ligands or other residues with the same charge. The torsion angles for this residue are unusual. Only glycine is flexible enough to make these torsion angles, mutation into another residue will force the local backbone into an incorrect conformation and will disturb the local structure.

**Figure 8:**
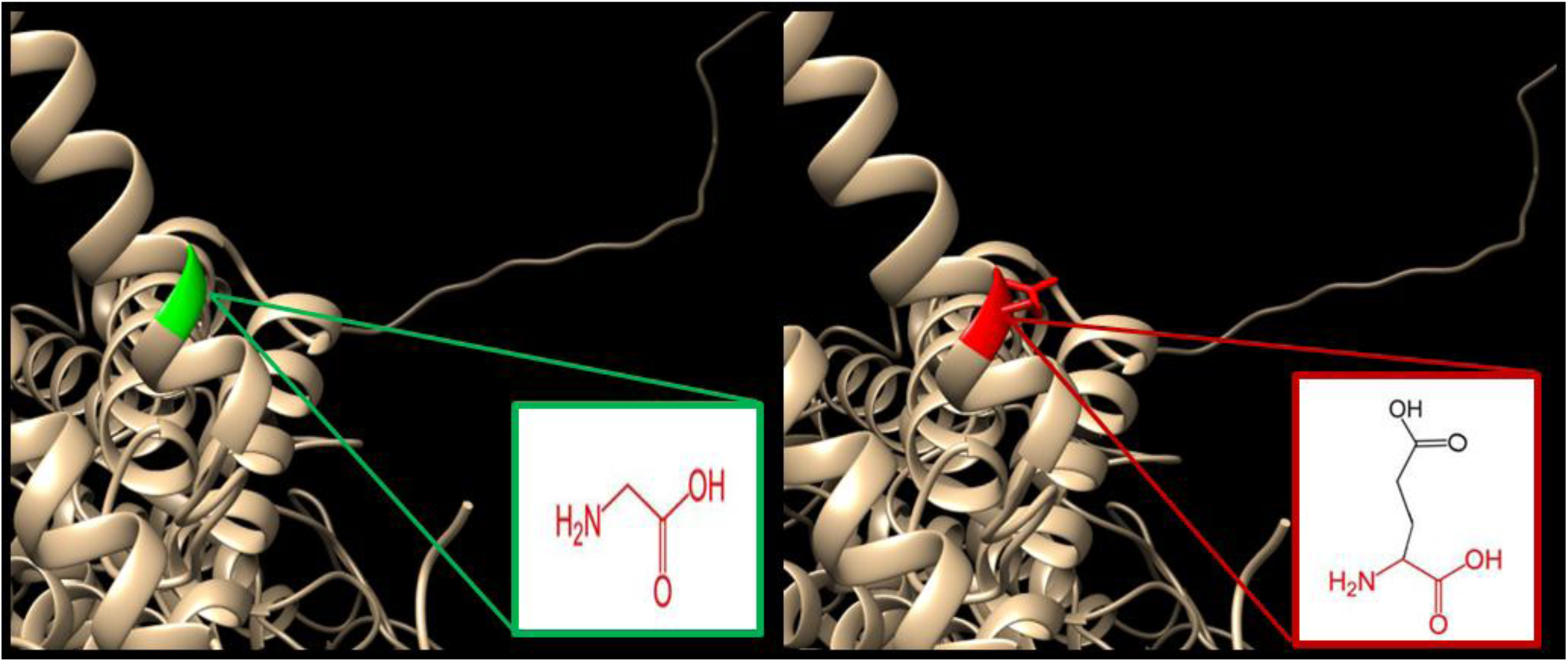
(G504E): The amino acid Glycine changes to Glutamate at position 495.

We also used Consurf server to check the conservation region of ATL1 protein, the result shows 4 SNPs (V67F, T120I, R217Q, and R495W) located in highly conserved regions, which can directly affect the protein function. **(Figure 9)**

**Figure 9:**
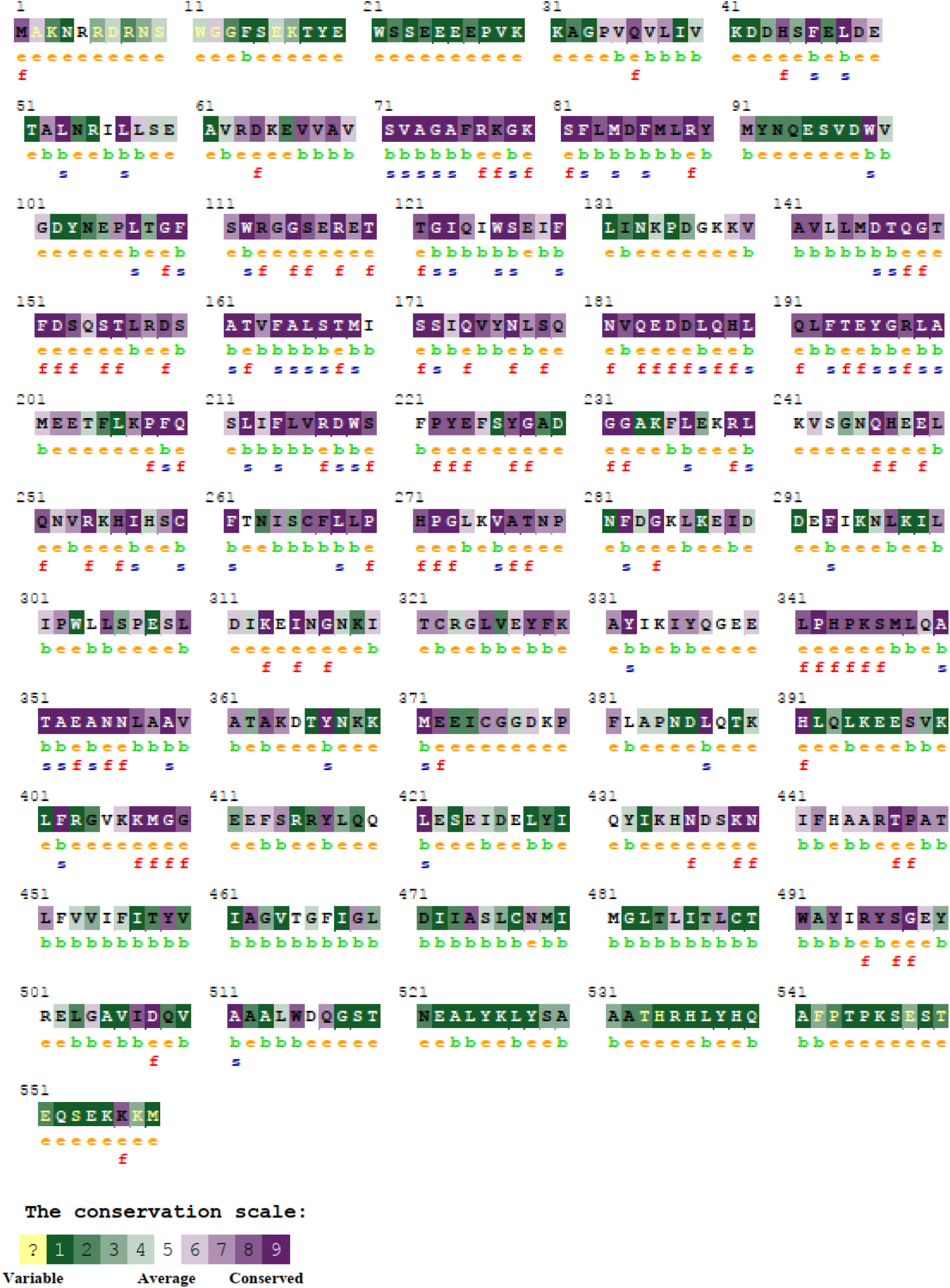
The conserved amino acids across species in CEBPA protein were determined using ConSurf. e: exposed residues according to the neural-network algorithm are indicated in orange letters. b: residues predicted to be buried are demonstrated via green letters. f: predicted functional residues (highly conserved and exposed) are indicated with red letters. s: predicted structural residues (highly conserved and buried) are demonstrated in blue letters. I: insufficient data (the calculation for this site performed in less than 10% of the sequences) is demonstrated in yellow letters.

We also used GeneMANIA which shown that *ATL1* has many dynamic functions: endomembrane system organization, endoplasmic reticulum organization, protein homooligomerization. The genes co-expressed with, sharing similar protein domain, or contributed to achieve similar function are shown in (Tables 4 & 5; Figure 10)

**Table 4:** The *ALT1* gene functions and its appearance in network and genome:

| Function | FDR | Genes in network | Genes in genome |
| --- | --- | --- | --- |
| endoplasmic reticulum organization | 1.91E-07 | 5 | 19 |
| endomembrane system organization | 0.0006584 | 6 | 210 |
| Golgi organization | 0.0956682 | 3 | 45 |
| cellular response to interferon-gamma | 0.5758816 | 3 | 90 |
| response to interferon-gamma | 0.7482479 | 3 | 106 |
| protein homooligomerization | 0.8135839 | 3 | 116 |

**Abbreviation: FDR:** false discovery rate, it is greater than or equal to the probability that this is a false positive.

**Table 5:** The gene co-expression, shared domain, and interaction with *ALT1* gene network:

| Gene 1 | Gene 2 | Weight | Network group |
| --- | --- | --- | --- |
| *GBP2* | *GBP1* | 0.015766 | Co-expression |
| *GBP2* | *GBP1* | 0.012037 | Co-expression |
| *GBP5* | *GBP4* | 0.007623 | Co-expression |
| *CACNA2D3* | *ATL1* | 0.010725 | Co-expression |
| *ATP8A2* | *ATL1* | 0.012415 | Co-expression |
| *KIAA0319* | *ATL1* | 0.015208 | Co-expression |
| *GBP2* | *GBP1* | 0.012014 | Co-expression |
| *GBP2* | *GBP1* | 0.022107 | Co-expression |
| *RTN4* | *GBP1* | 0.022003 | Co-expression |
| *GBP2* | *GBP1* | 0.007599 | Co-expression |
| *GBP1* | *GBP5* | 0.023868 | Co-expression |
| *GBP2* | *GBP3* | 0.027732 | Co-expression |
| *GBP2* | *GBP5* | 0.033093 | Co-expression |
| *GBP2* | *GBP1* | 0.031466 | Co-expression |
| *GBP2* | *GBP1* | 0.007406 | Co-expression |
| *GBP4* | *GBP3* | 0.021732 | Co-expression |
| *GBP5* | *GBP3* | 0.009855 | Co-expression |
| *GBP5* | *GBP4* | 0.010083 | Co-expression |
| *CACNA2D3* | *ATL1* | 0.03003 | Co-expression |
| *RTN4* | *GBP7* | 0.002513 | Co-expression |
| *RNFT2* | *ATL1* | 0.025588 | Co-expression |
| *SFXN3* | *ATL1* | 0.029366 | Co-expression |
| *TMOD2* | *ATL1* | 0.022688 | Co-expression |
| *GBP2* | *GBP1* | 0.014874 | Co-expression |
| *GBP5* | *GBP4* | 0.009542 | Co-expression |
| *GBP1* | *GBP3* | 0.018849 | Co-expression |
| *GBP1* | *GBP5* | 0.006281 | Co-expression |
| *GBP2* | *GBP3* | 0.015121 | Co-expression |
| *GBP2* | *GBP4* | 0.0151 | Co-expression |
| *GBP2* | *GBP5* | 0.006252 | Co-expression |
| *GBP2* | *GBP1* | 0.008656 | Co-expression |
| *GBP4* | *GBP3* | 0.014169 | Co-expression |
| *GBP5* | *GBP4* | 0.016914 | Co-expression |
| *GBP1* | *GBP3* | 0.016417 | Co-expression |
| *GBP1* | *GBP4* | 0.021962 | Co-expression |
| *GBP2* | *GBP3* | 0.009148 | Co-expression |
| *GBP2* | *GBP1* | 0.013121 | Co-expression |
| *RTN4* | *ATL1* | 0.008207 | Co-expression |
| *ATP8A2* | *ATL1* | 0.005958 | Co-expression |
| *SFXN3* | *RNF112* | 0.008297 | Co-expression |
| *TMOD2* | *ATL1* | 0.006243 | Co-expression |
| *KIAA0319* | *ATL1* | 0.003415 | Co-expression |
| *KIAA0319* | *RNFT2* | 0.013708 | Co-expression |
| *KIAA0319* | *ATP8A2* | 0.009184 | Co-expression |
| *RTN3* | *ATL1* | 0.002368 | Co-expression |
| *RTN3* | *RTN4* | 0.008981 | Co-expression |
| *RTN3* | *TMOD2* | 0.006031 | Co-expression |
| *GBP2* | *GBP1* | 0.003804 | Co-expression |
| *GBP2* | *GBP1* | 0.016589 | Co-expression |
| *GBP2* | *GBP1* | 0.021879 | Co-expression |
| *KIAA0319* | *ATP8A2* | 0.018812 | Co-expression |
| *GBP2* | *GBP1* | 0.007589 | Co-expression |
| *GBP1* | *GBP4* | 0.008522 | Co-localization |
| *GBP2* | *GBP1* | 0.003213 | Co-localization |
| *CACNA2D3* | *ATL1* | 0.003932 | Co-localization |
| *RTN4* | *ATL1* | 0.003111 | Co-localization |
| *RNFT2* | *ATL1* | 0.005352 | Co-localization |
| *ATP8A2* | *ATL1* | 0.005232 | Co-localization |
| *KIAA0319* | *ATL1* | 0.004243 | Co-localization |
| *ATL2* | *ATL3* | 0.018569 | Physical Interactions |
| *RTN4* | *ATL3* | 0.2318 | Physical Interactions |
| *RTN3* | *ATL3* | 0.2318 | Physical Interactions |
| *RTN4* | *ATL3* | 0.121214 | Physical Interactions |
| *RTN4* | *ATL1* | 0.113992 | Physical Interactions |
| *TMED9* | *ATL1* | 0.24926 | Physical Interactions |
| *RTN3* | *ATL1* | 0.158274 | Physical Interactions |
| *RTN3* | *ATL2* | 0.34741 | Physical Interactions |
| *RTN3* | *RTN4* | 0.036872 | Physical Interactions |
| *RTN3* | *RTN4* | 0.054499 | Physical Interactions |
| *GBP2* | *GBP1* | 1 | Predicted |
| *MUL1* | *ATL1* | 0.707107 | Predicted |
| *ATL2* | *MUL1* | 0.707107 | Predicted |
| *RTN4* | *ATL1* | 0.979022 | Predicted |
| *GBP7* | *ATL1* | 0.088508 | Shared protein domains |
| *GBP6* | *ATL1* | 0.088508 | Shared protein domains |
| *GBP6* | *GBP7* | 0.088508 | Shared protein domains |
| *GBP3* | *ATL1* | 0.088508 | Shared protein domains |
| *GBP3* | *GBP7* | 0.088508 | Shared protein domains |
| *GBP3* | *GBP6* | 0.088508 | Shared protein domains |
| *GBP4* | *ATL1* | 0.088508 | Shared protein domains |
| *GBP4* | *GBP7* | 0.088508 | Shared protein domains |
| *GBP4* | *GBP6* | 0.088508 | Shared protein domains |
| *GBP4* | *GBP3* | 0.088508 | Shared protein domains |
| *ATL3* | *ATL1* | 0.088508 | Shared protein domains |
| *ATL3* | *GBP7* | 0.088508 | Shared protein domains |
| *ATL3* | *GBP6* | 0.088508 | Shared protein domains |
| *ATL3* | *GBP3* | 0.088508 | Shared protein domains |
| *ATL3* | *GBP4* | 0.088508 | Shared protein domains |
| *GBP5* | *ATL1* | 0.088508 | Shared protein domains |
| *GBP5* | *GBP7* | 0.088508 | Shared protein domains |
| *GBP5* | *GBP6* | 0.088508 | Shared protein domains |
| *GBP5* | *GBP3* | 0.088508 | Shared protein domains |
| *GBP5* | *GBP4* | 0.088508 | Shared protein domains |
| *GBP5* | *ATL3* | 0.088508 | Shared protein domains |
| *ATL2* | *ATL1* | 0.088508 | Shared protein domains |
| *ATL2* | *GBP7* | 0.088508 | Shared protein domains |
| *ATL2* | *GBP6* | 0.088508 | Shared protein domains |
| *ATL2* | *GBP3* | 0.088508 | Shared protein domains |
| *ATL2* | *GBP4* | 0.088508 | Shared protein domains |
| *ATL2* | *ATL3* | 0.088508 | Shared protein domains |
| *ATL2* | *GBP5* | 0.088508 | Shared protein domains |
| *GBP1* | *ATL1* | 0.088508 | Shared protein domains |
| *GBP1* | *GBP7* | 0.088508 | Shared protein domains |
| *GBP1* | *GBP6* | 0.088508 | Shared protein domains |
| *GBP1* | *GBP3* | 0.088508 | Shared protein domains |
| *GBP1* | *GBP4* | 0.088508 | Shared protein domains |
| *GBP1* | *ATL3* | 0.088508 | Shared protein domains |
| *GBP1* | *GBP5* | 0.088508 | Shared protein domains |
| *GBP1* | *ATL2* | 0.088508 | Shared protein domains |
| *GBP2* | *ATL1* | 0.088508 | Shared protein domains |
| *GBP2* | *GBP7* | 0.088508 | Shared protein domains |
| *GBP2* | *GBP6* | 0.088508 | Shared protein domains |
| *GBP2* | *GBP3* | 0.088508 | Shared protein domains |
| *GBP2* | *GBP4* | 0.088508 | Shared protein domains |
| *GBP2* | *ATL3* | 0.088508 | Shared protein domains |
| *GBP2* | *GBP5* | 0.088508 | Shared protein domains |
| *GBP2* | *ATL2* | 0.088508 | Shared protein domains |
| *GBP2* | *GBP1* | 0.088508 | Shared protein domains |
| *RNF112* | *ATL1* | 0.036836 | Shared protein domains |
| *RNF112* | *GBP7* | 0.036836 | Shared protein domains |
| *RNF112* | *GBP6* | 0.036836 | Shared protein domains |
| *RNF112* | *GBP3* | 0.036836 | Shared protein domains |
| *RNF112* | *GBP4* | 0.036836 | Shared protein domains |
| *RNF112* | *ATL3* | 0.036836 | Shared protein domains |
| *RNF112* | *GBP5* | 0.036836 | Shared protein domains |
| *RNF112* | *ATL2* | 0.036836 | Shared protein domains |
| *RNF112* | *GBP1* | 0.036836 | Shared protein domains |
| *RNF112* | *GBP2* | 0.036836 | Shared protein domains |
| *RTN3* | *RTN4* | 0.333333 | Shared protein domains |
| *GBP7* | *ATL1* | 0.086491 | Shared protein domains |
| *GBP6* | *ATL1* | 0.086491 | Shared protein domains |
| *GBP6* | *GBP7* | 0.11064 | Shared protein domains |
| *GBP3* | *ATL1* | 0.086491 | Shared protein domains |
| *GBP3* | *GBP7* | 0.11064 | Shared protein domains |
| *GBP3* | *GBP6* | 0.11064 | Shared protein domains |
| *GBP4* | *ATL1* | 0.086491 | Shared protein domains |
| *GBP4* | *GBP7* | 0.11064 | Shared protein domains |
| *GBP4* | *GBP6* | 0.11064 | Shared protein domains |
| *GBP4* | *GBP3* | 0.11064 | Shared protein domains |
| *ATL3* | *ATL1* | 0.086491 | Shared protein domains |
| *ATL3* | *GBP7* | 0.11064 | Shared protein domains |
| *ATL3* | *GBP6* | 0.11064 | Shared protein domains |
| *ATL3* | *GBP3* | 0.11064 | Shared protein domains |
| *ATL3* | *GBP4* | 0.11064 | Shared protein domains |
| *GBP5* | *ATL1* | 0.086491 | Shared protein domains |
| *GBP5* | *GBP7* | 0.11064 | Shared protein domains |
| *GBP5* | *GBP6* | 0.11064 | Shared protein domains |
| *GBP5* | *GBP3* | 0.11064 | Shared protein domains |
| *GBP5* | *GBP4* | 0.11064 | Shared protein domains |
| *GBP5* | *ATL3* | 0.11064 | Shared protein domains |
| *ATL2* | *ATL1* | 0.086491 | Shared protein domains |
| *ATL2* | *GBP7* | 0.11064 | Shared protein domains |
| *ATL2* | *GBP6* | 0.11064 | Shared protein domains |
| *ATL2* | *GBP3* | 0.11064 | Shared protein domains |
| *ATL2* | *GBP4* | 0.11064 | Shared protein domains |
| *ATL2* | *ATL3* | 0.11064 | Shared protein domains |
| *ATL2* | *GBP5* | 0.11064 | Shared protein domains |
| *GBP1* | *ATL1* | 0.086491 | Shared protein domains |
| *GBP1* | *GBP7* | 0.11064 | Shared protein domains |
| *GBP1* | *GBP6* | 0.11064 | Shared protein domains |
| *GBP1* | *GBP3* | 0.11064 | Shared protein domains |
| *GBP1* | *GBP4* | 0.11064 | Shared protein domains |
| *GBP1* | *ATL3* | 0.11064 | Shared protein domains |
| *GBP1* | *GBP5* | 0.11064 | Shared protein domains |
| *GBP1* | *ATL2* | 0.11064 | Shared protein domains |
| *GBP2* | *ATL1* | 0.086491 | Shared protein domains |
| *GBP2* | *GBP7* | 0.11064 | Shared protein domains |
| *GBP2* | *GBP6* | 0.11064 | Shared protein domains |
| *GBP2* | *GBP3* | 0.11064 | Shared protein domains |
| *GBP2* | *GBP4* | 0.11064 | Shared protein domains |
| *GBP2* | *ATL3* | 0.11064 | Shared protein domains |
| *GBP2* | *GBP5* | 0.11064 | Shared protein domains |
| *GBP2* | *ATL2* | 0.11064 | Shared protein domains |
| *GBP2* | *GBP1* | 0.11064 | Shared protein domains |
| *RNF112* | *ATL1* | 0.063936 | Shared protein domains |
| *RNF112* | *GBP7* | 0.039812 | Shared protein domains |
| *RNF112* | *GBP6* | 0.039812 | Shared protein domains |
| *RNF112* | *GBP3* | 0.039812 | Shared protein domains |
| *RNF112* | *GBP4* | 0.039812 | Shared protein domains |
| *RNF112* | *ATL3* | 0.039812 | Shared protein domains |
| *RNF112* | *GBP5* | 0.039812 | Shared protein domains |
| *RNF112* | *ATL2* | 0.039812 | Shared protein domains |
| *RNF112* | *GBP1* | 0.039812 | Shared protein domains |
| *RNF112* | *GBP2* | 0.039812 | Shared protein domains |
| *RTN3* | *RTN4* | 0.333333 | Shared protein domains |

**Figure 10:**
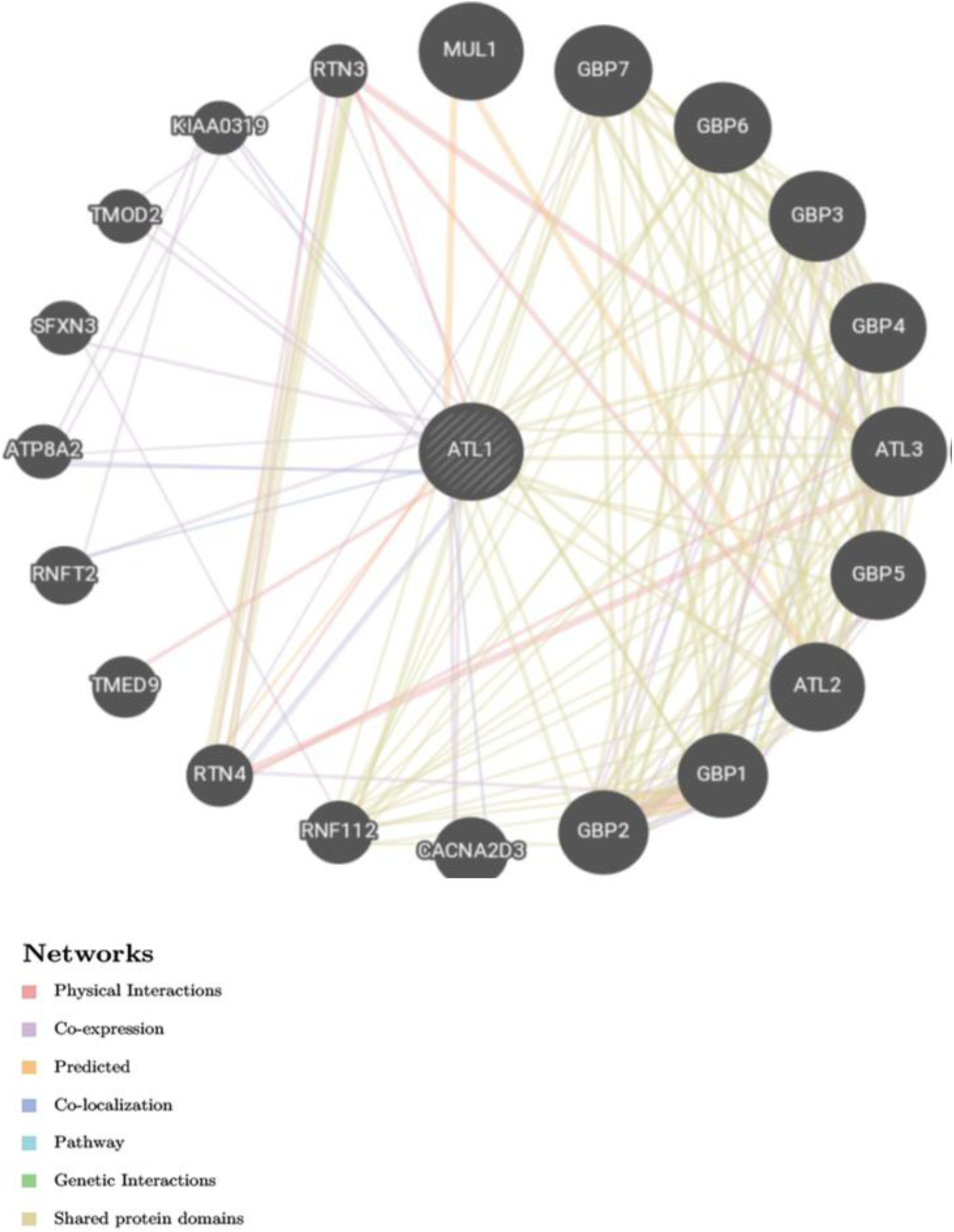
Biological network Interaction between *ATL1* and its related genes.

We also used ClinVar to compare our results that had been found by in silico approach with the clinical one, in (R217Q) SNP was found to be pathogenic, our result doesn’t matches with this result, [48] however, some evidence-based studies matches with our result.[49, 50] The second SNP (R495W) the associated clinical studies in ClinVar show that our result matches with the reported record which is pathogenic variant.[51] While for the other SNPs (V67F, T120I & G504E) we didn’t found any associated clinical studies.

The Variant Effect Predictor annotates mutations using an extensive array of reference data from previously detected mutations, evidence based results, and estimation of biophysical consequences of mutations; and that is what makes VEP an accurate web-based tool.[42] VEP described regulatory consequences for several mutations, including 15 mutations within a coding region, 15 mutations within a non-coding region, 2 mutations within upstream gene, 9 mutations within downstream gene, 1 mutation within non-coding transcript exon, and 2 mutations within 5 prime UTR variant; briefly, mutations within a coding region affect the protein function, while mutations within non-coding regions can significantly affect disease and could be contribute in the phenotypic feature and RNA-binding proteins (RBPs);[52, 53] while mutations in the upstream, downstream, 5′-, and 3′-UTRs might affect transcription or translation process.[54] The consequences are shown in (Table 6) while (Figure 2) demonstrates the summary pie charts and statistics.

**Table 6:**
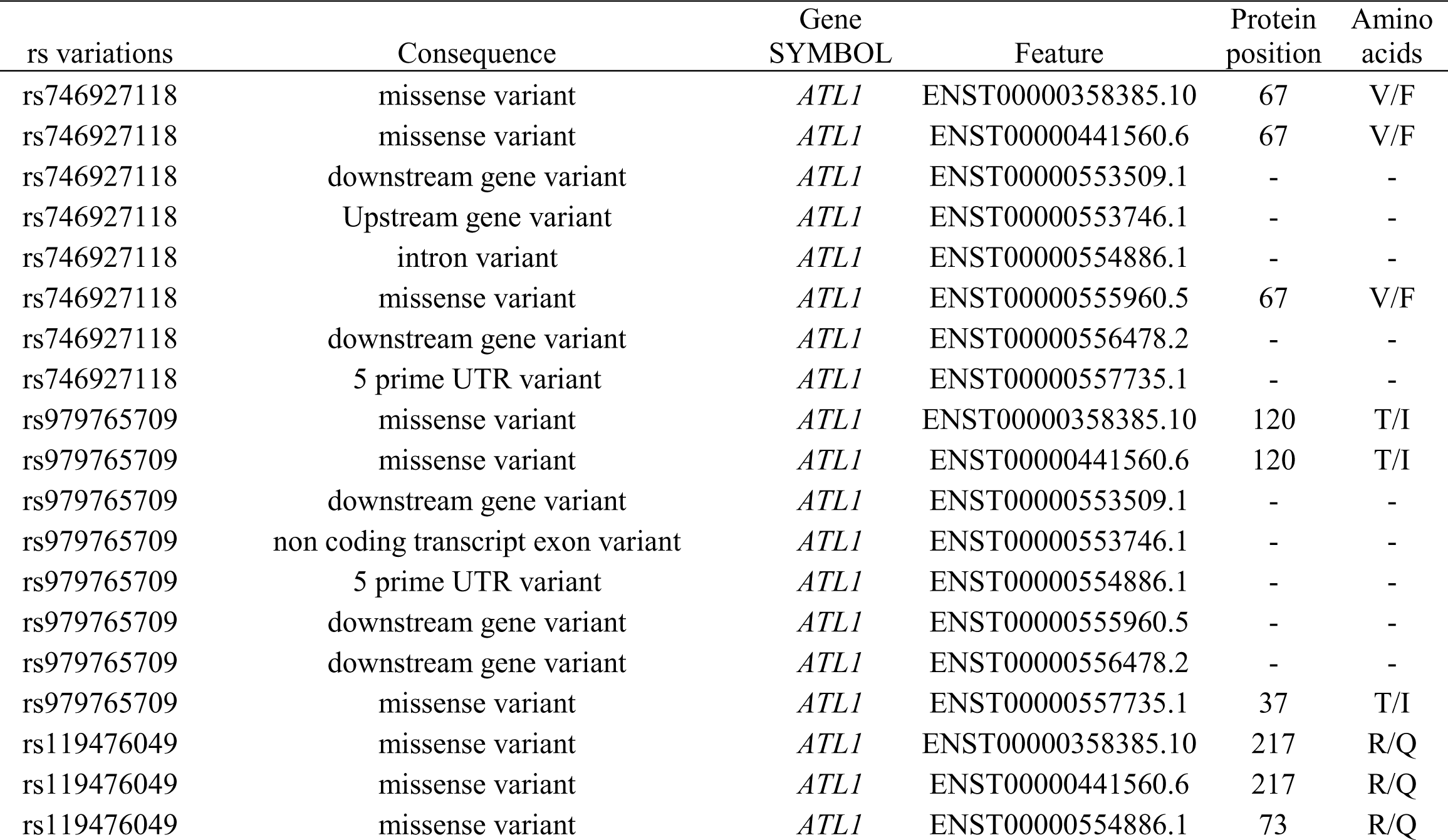
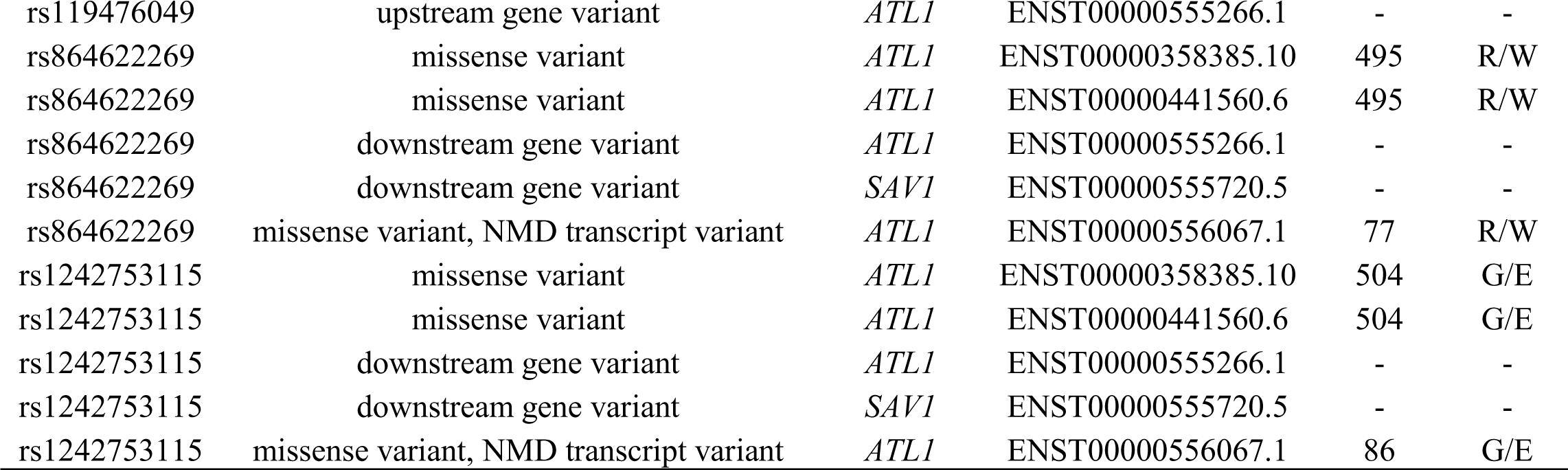
Shows variant consequences, transcripts, and regulatory features by VEP tool:

This study is the first bioinformatics analysis while all other studies were in vivo and in vitro analysis.[17, 18] To conclude, 5 disease-causing mutations were recognized as the most pathogenic SNPs in the coding region of *ATL1* gene that may cause SPG3A, and therefore, it may be used as genomic biomarkers for SPG3A. Lastly, Wet lab techniques are suggested to backing these outcomes.

## 5. Conclusion

In this study the impact of nsSNPs in the *ATL1 gene* was investigated by various bioinformatics tools, that revealed presence of five deleterious SNPs (V67F, T120I, R217Q, R495W and G504E), which have a functional impact on ATL1 protein; and therefore, can be used as genomic biomarkers specifically before 4 years old; also it may play a key role in pharmacogenomics by evaluating drug response for this disabling disease.

## Conflicts of Interest

The authors declare that there are no conflicts of interest regarding the publication of this paper.

## Data Availability

All data underlying the results are available as part of the article, and no additional source data are required.

## Funding

The authors received no financial support for the research, authorship, and/or publication of this article.

